# Tbx1 haploinsufficiency causes brain metabolic and behavioral anomalies in adult mice which are corrected by vitamin B12 treatment

**DOI:** 10.1101/2024.02.01.578212

**Authors:** Marianna Caterino, Debora Paris, Giulia Torromino, Michele Costanzo, Gemma Flore, Annabella Tramice, Elisabetta Golini, Silvia Mandillo, Diletta Cavezza, Claudia Angelini, Margherita Ruoppolo, Andrea Motta, Elvira De Leonibus, Antonio Baldini, Elizabeth Illingworth, Gabriella Lania

## Abstract

**Introduction:** The brain-related phenotypes observed in 22q11.2 deletion syndrome (22q11.2DS) are highly variable and their origin is poorly understood. Changes in brain metabolism may cause or contribute to the phenotypes, given that many of the deleted genes (approx. 10%) are implicated in metabolic processes, but this is currently unknown. It is clearly important to address this knowledge gap, but in humans, the primary material required for studying brain metabolism is inaccessible. For this reason, we sought to address the issue using two mouse models of 22q11.2DS.

**Methods:** We used three independent approaches to investigate brain metabolism in young adult mice, namely, mass spectrometry, nuclear magnetic resonance spectroscopy and transcriptomics. We selected to study primarily *Tbx1* single gene mutants because it is the primary candidate disease gene. We then confirmed key findings in the multi-gene deletion mutant *Df1/+*.

**Results:** We found that *Tbx1* mutants have alterations of specific brain metabolites, including methylmalonic acid, which is highly brain-toxic, as well as a more general metabolomic imbalance. We provide transcriptomic evidence of an interaction genotype-vB12 treatment, and behavioural evidence of a response to vB12 treatment, which rescued some of the behavioural anomaly observed in *Tbx1* mutants. We conclude that *Tbx1* haploinsufficiency causes extensive brain metabolic anomalies, which are partially responsive to vB12 treatment. We suggest that alterations of glutamine-glutamate metabolism and fatty acid metabolism are key components of the metabolic phenotype in these mutants.

## Introduction

22q11.2 deletion syndrome is associated with a broad range of brain-related clinical signs and symptoms that is highly variable between patients, even within the same family. This includes structural brain anomalies, motor and cognitive deficits, learning difficulties, and a greatly increased risk for psychiatric disorders, especially anxiety and schizophrenia (McDonald-McGinn et al., 2015). The contribution of individual deleted genes to the brain phenotypes is largely unknown, or unproven, due to the rarity of point mutations. However, rare point mutations indicate that *TBX1* haploinsufficiency is responsible for most of the physical abnormalities associated with 22q11.2DS, and they have also been associated with attention deficits, mild mental retardation, learning difficulties, developmental delay, Asperger syndrome and depression (Paylor et al., 2006)(Torres-Juan et al., 2007)(Ogata et al., 2014)(Haddad et al., 2019). However, the different nature of the brain phenotypes found in 22q11.2DS patients suggests that additional genes from the deleted region likely play a role.

Intriguingly, clinical studies suggest that 22q11.2DS patients might have altered brain metabolism (Napoli et al., 2015)(Korteling et al., 2022)(Zafarullah et al., 2024). In particular, a study of the metabolome and of mitochondrial function in a small group of children with 22q11.2DS revealed a distinct biochemical profile, which was consistent with increased oxidative stress, and shared features with congenital propionic and/or methylmalonic acidemia (Napoli et al., 2015).

Given the high genetic and phenotypic relevance of the available mouse models of 22q11.2DS, they provide, potentially, a unique opportunity to corroborate newly identified phenotypes in patients and to study the pathogenetic mechanisms underlying the brain-related deficits. In support of this notion, is the finding that many orthologs of the del22q11.2 genes are expressed in the mouse brain (Maynard et al., 2003), and at least nine orthologs are involved in mitochondrial metabolism (Maynard et al., 2008)(Meechan et al., 2011)(Devaraju and Zakharenko, 2017). Thus, combined heterozygosity of multiple mitochondrial genes might negatively impact brain development and/or brain function, especially given high energy demands of the brain.

We reasoned that mouse models of 22q11.2DS might have metabolic disturbances similar to those reported in patients, in which case, *in vivo* genetic and pharmacological manipulation of the model should provide insights into the contribution of the metabolic phenotypes to disease pathogenesis and/or mechanisms. Indeed, we have previously shown that vitamin B12 (vB12, cyanocobalamin) treatment ameliorates some of the heart and brain abnormalities observed in *Tbx1* mutant mice (Lania et al., 2016)(Favicchia et al., 2021)(Lania et al., 2022), concluding that vB12 acts downstream of *Tbx1* because it rescues anatomical brain anomalies in *Tbx1* homozygous mutants during embryonic and adult life (Favicchia et al., 2021). Vitamin B12 acts as a cofactor in the methionine cycle for the production of reaction intermediates such as SAM (S-Adenosyl methionine), which in turn provides methyl groups for the methylation of proteins and nucleic acids. In addition, in the mitochondria, it is a cofactor of Methyl malonyl-CoA mutase, the enzyme that catalyses the reversible isomerization of Methyl malonyl-CoA to Succinyl-CoA, an essential intermediate in the citric acid cycle (TCA, Krebs cycle). Alterations of this pathway can lead to an accumulation of methylmalonic acid (MMA), which is highly brain toxic.

In this study, we applied a multidisciplinary approach involving metabolomic, transcriptional and behavioural studies to search for metabolic alterations in the brain of adult *Tbx1* heterozygous mice. Through this approach, we identified a new metabolic phenotype that was associated with reduced sensorimotor gating deficits in *Tbx1^+/-^* mice. We also discovered that the metabolic and behavioural phenotype responded to vB12 treatment. In particular, the biochemical analysis revealed the accumulation of methylmalonic acid (MMA) and alteration of metabolites in vB12-related pathways in the brain of adult *Tbx1^+/-^* mice, and in mice carrying a multigenic deletion (*Df1/+*) of 22q11.2DS orthologs that includes *Tbx1* (Lindsay et al., 1999).

Overall, our results show that *Tbx1* haploinsufficiency is associated with significant metabolic abnormalities in the young adult brain. We found that post-natal vB12 treatment corrects some of the metabolic alterations and eradicates the PPI anomaly.

## Results

### Methylmalonic acid (MMA) accumulates in the brain of Tbx1 and Df1 heterozygous mice

We were intrigued by the possibility that patients with 22q11.2DS have altered levels of brain toxic metabolites related to the vB12 pathway. We determined whether this was the case in the mouse models. For this, we first performed a preliminary metabolome analysis using isolated whole brains of male and female *Tbx1^+/-^* (n= 18) and WT (n= 10) mice between one and two months of age. A set of metabolites was quantified in brain extracts by liquid chromatography-tandem mass spectrometry (LC-MS/MS) (Supplementary Table 1). The significant quantitative metabolic alterations in *Tbx1^+/-^*mice were provided by univariate analysis, as shown in the volcano plot (Figure 1A). The metabolites showing significant differences in *Tbx1^+/-^ vs* WT were MMA, which was two-fold higher in *Tbx1^+/-^* brains compared to WT brains, LA (Lactic Acid, C10DC (Decanedioylcarnitine), C18:1 (Octadecenoylcarnitine) and Asp (Aspartic Acid) (Figure 1A).

**Figure 1.**
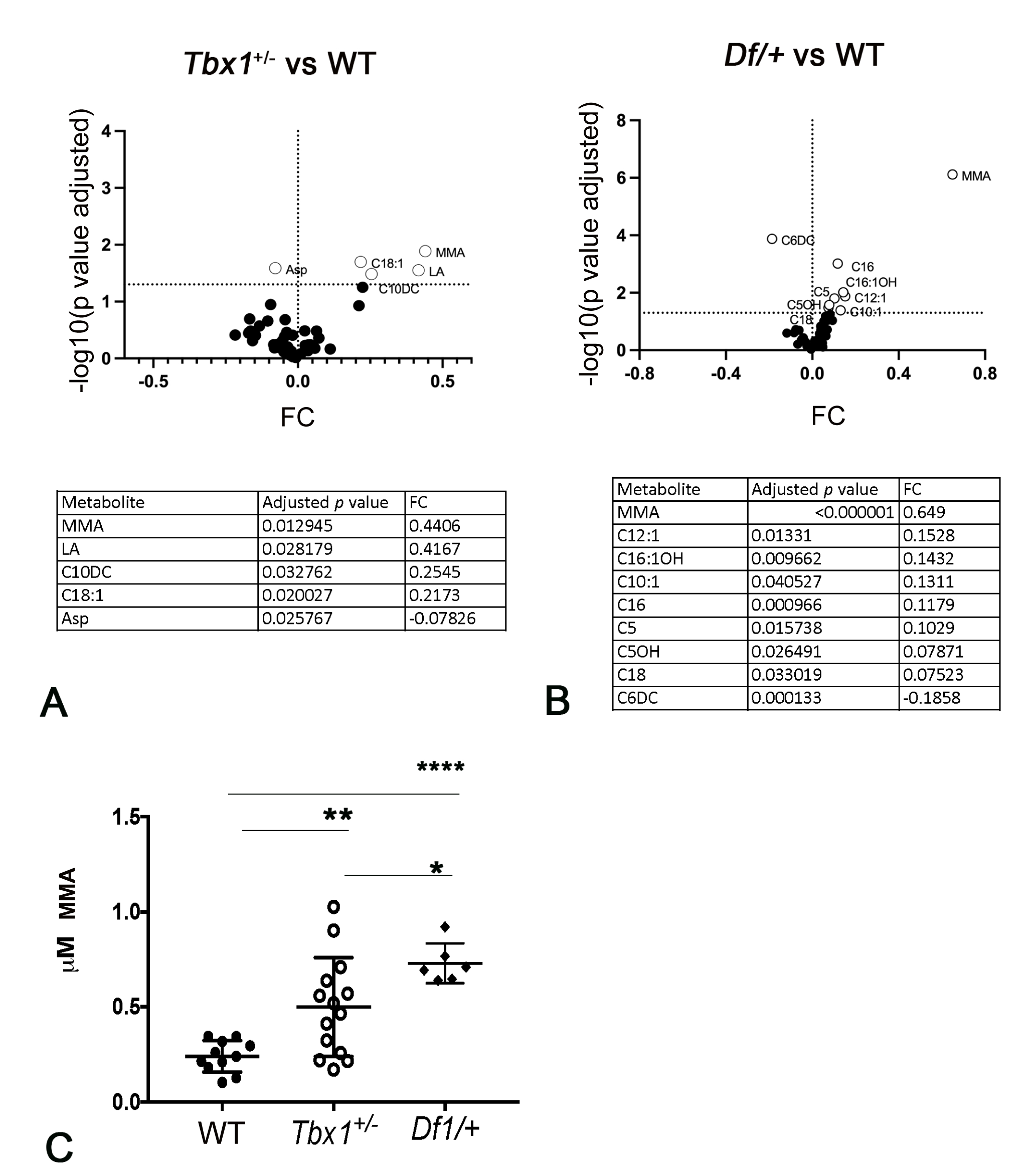
Targeted metabolome analysis by liquid chromatography - tandem mass spectrometry (LC-MS/MS). Volcano plots showing differential concentrations of selected metabolites in A) *Tbx1^+/-^* vs WT and B) *Df1/+* vs WT brains. The white dots represent the differential metabolites and the black dots all the metabolites identified in the dataset, the relative abundance of which was not significantly different between the groups. The differential metabolites are listed in the tables accompanied by their corresponding values of difference (FC, fold change) and adjusted p-values. C) Abundance of MMA (µM, mean ± SEM) in WT, *Tbx1^+/-^* and *Df1/+* brains. Differences between groups were evaluated by performing the ordinary one-way ANOVA test and Holm-Sidak’s multiple comparison test (* p<0.05, ** p<0.01, **** p<0.0001). Each symbol in the plot represents an individual animal.

The human deletion is better modelled by *Df1/+* mice, which carry a heterozygous deletion of 27 orthologs of del22q11.2 genes, 9 of which encode metabolic enzymes. We analysed whole brain tissue isolated from *Df1/+* and WT (control) mice (n = 5 per genotype). Results showed that the mean concentration of MMA was three-fold higher in *Df1/+* brains compared to WT brains (Figure 1B, 1C). In addition, C12:1 (Dodecenoylcarnitine), C16:1OH (3-Hydroxyhexadecenoylcarnitine), C10:1 (Decenoylcarnitine), C16 (Hexadecanoylcarnitine), C5 (Valerylcarnitine), C5OH (3-Hydroxyisovalerylcarnitine/3-hydroxy-2 methylbutyrylcarnitine), C18 (Octadecanoylcarnitine) were increased in the *Df1/+* brain; conversely, C6DC (Methylglutarylcarnitine) was reduced (Figure 1B).

Interestingly, we did not find any changes in MMA concentration in the brain of pre-term *Tbx1^+/-^* embryos, nor in non-CNS tissues of adult male and female *Tbx1^+/-^* mice (Supplementary Figure 1), suggesting that the phenotype is age- and tissue-dependent. These results suggest that *Tbx1* haploinsufficiency compromises mitochondrial metabolic processes related to vB12 (Figure 1C).

### Tbx1 haploinsufficiency alters the brain metabolomic profile of young adult mice

In order to visualise the effects of *Tbx1* haploinsufficiency on cell metabolism across the genome, we evaluated the metabolomic and transcriptional profiles of adult *Tbx1*^+/-^ and WT brains (n = 6 - 8 for each genotype) that were vB12- or vehicle (PBS)-treated. For this, we surgically removed the brain and separated the two hemispheres, one of which was used for NMR analysis, while the other was used for whole genome transcriptomic analysis by RNA quanTiseq. Each brain hemisphere was treated separately for the NMR analysis and for RNA-seq without pooling samples and each dataset was treated as a biological replicate.

The effects of the *Tbx1* mutation on brain metabolism were analysed by reporting the results obtained in vehicle-treated *Tbx1*^+/-^ and WT brains, followed by those obtained in vB12 treated animals with the same genotypes. For the NMR studies, data from each hemisphere were obtained from 1D and 2D NMR experiments. Hydrophilic and lipophilic metabolites were assigned to spectral signals by comparing the observed chemical shift with published data (Fan, 1996) and/or online databases (Wishart et al., 2022). To obtain the biochemical information from the NMR data, the 1D metabolic profiles underwent multivariate data analysis and projection methods. We applied an unsupervised principal component analysis (PCA) to assess class (genotype and/or treatment) homogeneity and to identify outliers then we used a supervised OPLS-DA (Orthogonal partial least squares discriminant analysis) to evaluate class discrimination and highlight metabolic differences associated with *Tbx1* mutation. The statistical analysis of the hydrophilic fraction yielded a regression model with high-quality parameters [R^2^= 0.99 (goodness of fit), Q^2^=0.94 (power in goodness of prediction) (L. Eriksson, T. Byrne, E. Johnsson, J. Trygg and C. Vikstrom) and CV-ANOVA test, *p* value = 0.0006], and a clear class discrimination (Figure 2A). The WT class (black dots) was located at negative t[1] values, while the *Tbx1*^+/-^ class (empty dots) is located at positive values. The discrimination along the t[1] parallel component (y axis) accounted for the genotype separation, while the orthogonal component to[1] (x axis) accounted for the intra-class homogeneity (Figure 2A, 2A’, 2C). The associated S plot (Figure 2A’) indicated the NMR chemical shifts -then assigned to metabolites-that discriminated between the two classes..

**Figure 2.**
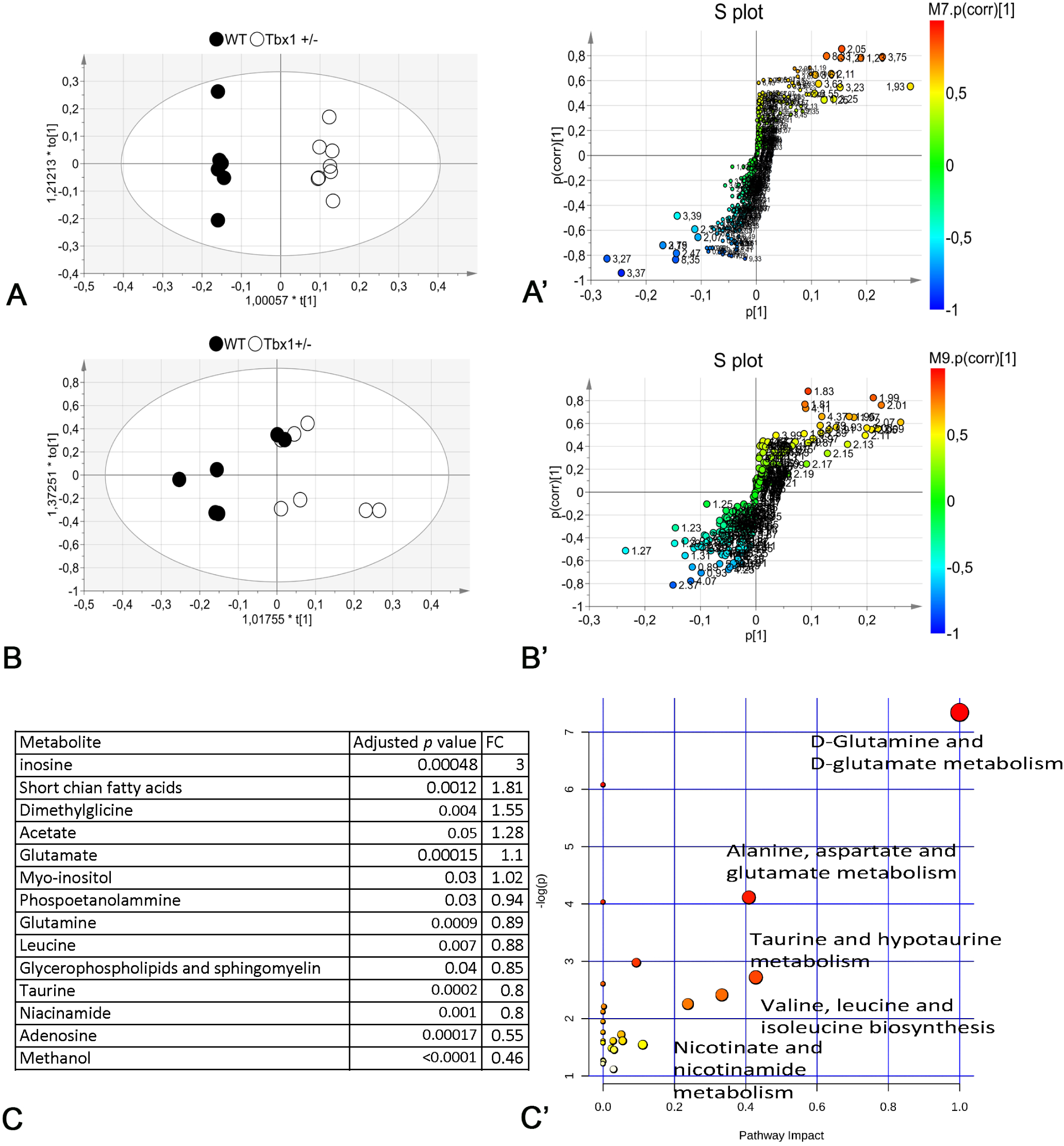
Brain metabolic profiles of *Tbx1 ^+/-^* and wild type brains. A-B) Scores plot obtained from the orthogonal partial least squares discriminant analysis (OPLS-DA) of brain extracts. NMR data from hydrophilic (A) and hydrophobic (B) phase. The principal components t[1] and to[1] describe the space in which the NMR metabolic profiles of each brain extract are projected. Spectra of WT and *Tbx1*^+/-^ are separated along the predictive t[1] axis. A’-B’) S-plot associated with the multivariate model providing principal component visualization to facilitate model interpretation. The NMR variables situated far out on the wings of the S in the left down corner present higher concentration in the wild-type class, while variables in the upper right side of the plot indicate metabolites with an increased concentration in Tbx1+/- mutant brain. The colour code refers to the correlation values. C) The differential metabolites are listed in the tables accompanied by their corresponding values of difference (FC, Fold changes) and adjusted p-value. C’ Discriminant metabolites were used as input for Pathway analysis in *Mus musculus* libraries to identify the most relevant pathways involved in the *Tbx1*^+/-^ mutant. Pathway impact on the axis represents a combination of the centrality and pathway enrichment results; higher impact values represent the relative importance of the pathway; the size of the circle indicates the impact of the pathway while the colour represents the significance (the more intense the red colour, the lower the p value).

The statistical analysis of the lipophilic fraction generated a weak OPLS-DA model with only limited class separation (with parameters R^2^= 0.56, Q^2^=0.28 and CV-ANOVA test *p* value =0.6), with a single metabolite responsible for the class distinction (Figure 2B, 2B’, 2C and Supplementary Figure 2). Together, the hydrophilic and lipophilic results revealed a group of compounds that characterised the brain metabolic differences between *Tbx1*^+/-^ and WT mice (Figure 2B, 2C). In particular, for the hydrophilic fraction, we found several metabolites to be significantly different between *Tbx1*^+/-^ and WT mice. Specifically, in *Tbx1*^+/-^ mice, we observed downregulation of adenosine, niacinamide (vitamin B3), methanol, phosphoethanolamine, glutamine, leucine, glycerophospholipids and sphingomyelin (from the lipophilic fraction) and taurine, and upregulation of inosine, short-chain fatty acids, acetate, glutamate, myoinositol and dimethylglycine (Figure 2C Supplementary Figure 2). To understand better the biological relevance of the data, we investigated the metabolic pathways in which the differently regulated molecular species are involved using MetaboAnalyst 4.0 software (https://www.metaboanalyst.ca/). The pathways found are depicted in Figure 2C’ which reports the impact of each pathway versus *p* values..

Using the discriminating metabolites and considering only the pathways with an impact >40%, we found glutamine and glutamate metabolism (*p* = 7.65 × 10^-4^, impact 100%), alanine, aspartate and glutamate metabolism (*p* = 1.91 × 10^-2^, impact 41%), and taurine and hypotaurine metabolism (*p* = 7.13 × 10^-2^, impact 42%), to be potentially altered in *Tbx1*^+/-^ brains.

### Tbx1 haploinsufficiency has no significant effect on the brain transcriptome

We next sought to identify the transcriptional changes associated with the metabolic alterations identified in *Tbx1*^+/-^ mice. We performed whole brain tissue RNA quanTiseq analysis using the controlateral brain hemispheres to those used for NMR, and as for NMR, we performed the sequencing on individual hemispheres without pooling samples. Results showed that there were very few differentially expressed genes in *Tbx1*^+/-^ vs WT brains, (n=22 out of 14535 expressed genes (Fig. 5 and Supplementary Tab. 2). This surprising result might be due to a dilution effect on the target tissue (brain endothelial cells); *Tbx1* is only expressed in a subset of brain vessels (approx. 1.5% of brain cells) in the adult mouse (Ximerakis et al., 2019).

### Vitamin B12 treatment strongly affects the brain metabolome and transcriptome, and partially compensates for the effects of Tbx1 haploinsufficiency

Given the ability of vB12 treatment to rescue diverse phenotypes caused by *Tbx1* haploinsufficiency (Lania et al., 2016)(Favicchia et al., 2021)(Lania et al., 2022), we asked whether this occurs through metabolic mechanisms that are transcriptionally regulated by *Tbx1* or by correcting the brain metabolic profile of *Tbx1*^+/-^ mice.

To investigate the involvement of the vB12 pathway in the metabolic phenotype of *Tbx1^+/-^* and *Df1/+* mice, we treated mice postnatally with vB12 or vehicle (PBS) every three days for 28 days, between the age of 4 and 8 weeks, after which the animals were sacrificed. We first measured MMA concentration by LC-MS/MS in brain extracts. Results showed that *Tbx1^+/-^* and *Df1/+* brains have increased concentrations of MMA compared to WT controls (all animals PBS-treated), and they demonstrated that vB12 treatment restored MMA to wild type levels in both of these mutants (Figure 3).

**Figure 3.**
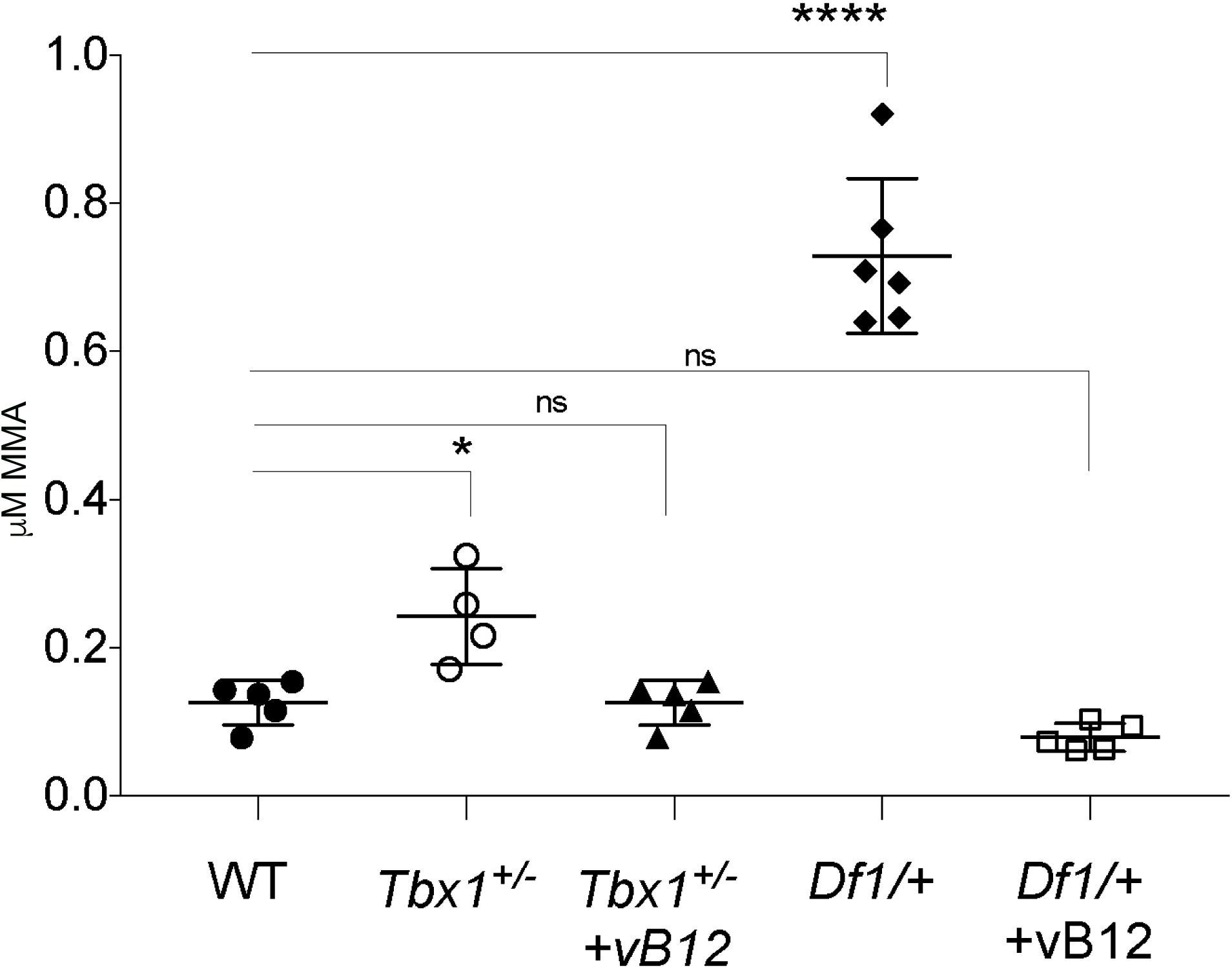
Vitamin B12 treatment restores MMA to wild type levels in *Tbx1^+/-^* and *Df1/+* mice. The abundance of MMA (µM, mean ± SEM) was evaluated in WT, *Tbx1^+/-^*, *Df1^/+^, Tbx1^+/-^* +vB12 and *Df1^/+^* +vB12. Differences between groups were evaluated by the ordinary one-way ANOVA test and the Holm-Sidak’s multiple comparison test (* p<0.05, ** p<0.01, **** p<0.0001). The normal distribution was verified according to D’Agostino and Pearson tests. Each symbol in the plot represents an individual animal.

Next, we evaluated the metabolomic and transcriptional profiles in a separate group of *Tbx1*^+/-^ and WT mice treated post-natally with vB12 or PBS, as described above. At the end of the treatment, we surgically removed the brains and performed NMR analysis and whole genome transcriptomic analysis on single brain hemispheres (n = 7 per genotype). In order to determine whether the metabolic changes observed in *Tbx1^+/-^* mice were related to the vB12 pathway, we applied NMR-based metabolomics analysis to the three classes (treatments and genotypes), namely, PBS-treated *Tbx1^+/-^* and WT mice and *Tbx1*^+/-^ mice treated with vB12. The OPLS-DA analysis yielded a regression model with high-quality parameters (R^2^ = 0.94, Q^2^ = 0.75 and CV-ANOVA test *p* value = 0.0008). The WT class (black dots) and the *Tbx1*^+/-^ class (empty dots) were both located at positive t[1] values, while the *Tbx1*^+/-^ vB12 class (empty black triangles) was located at negative values (Figure 4A). As can be observed from the scores plot, the first component t[1] (x axis) revealed a major alteration in the metabolic profile caused by the effect of vB12 treatment on *Tbx1*^+/-^ mice, evidenced by the distance between the vB12 treated *Tbx1*^+/-^ class (negative values) compared to the PBS-treated classes. In contrast, the second component t[2] (y axis) revealed only a minor alteration, due to the difference between PBS-treated *Tbx1^+/-^* and WT mice. By assigning metabolites to signals indicated in the associated loadings plot (Fig. 4B) and quantifying the variation in their content among classes, we observed that in only three cases did vB12 treatment rebalance a metabolic alteration observed in *Tbx1*^+/-^ mice, namely, inosine, glutamate and SCFAs (Fig. 4C). The ability of vB12 to rescue these particular metabolites, together with the known role of vB12 in the mitochondrial compartment, further suggest that the metabolic phenotype in *Tbx1*^+/-^ mice relates to mitochondrial activity.

**Figure 4.**
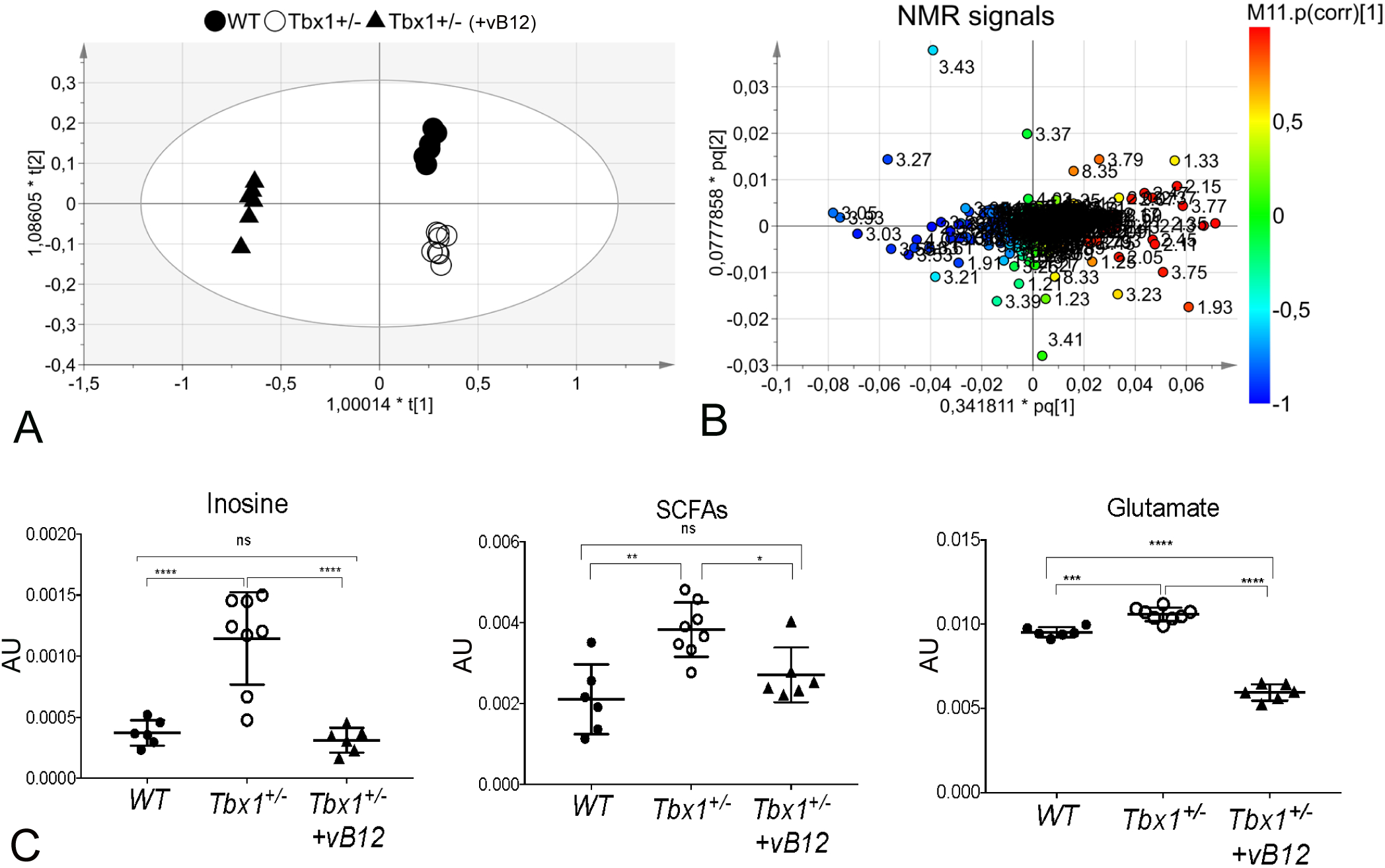
Metabolic profile of the *Tbx1^+/-^* brain after vitamin B12 treatment. The metabolic profiles of *Tbx1* mutant brain extracts were compared to those of wild type brains with and without vB12 treatment. A) Scores plot obtained from the orthogonal partial least squares’ discriminant analysis (OPLS-DA) of brain extracts (NMR data). B) Loadings plot associated to the scores plot in A) showing the NMR signals responsible for the data cluster. Each variable in the plot corresponds to a signal in the metabolic profile of the dataset. Combining scores and loadings plots by an ideal superposition, NMR variables located in the same area of samples in the scores plot are more expressed by that group while presenting a lower content in the class lying at the opposite site. C) Graphical representation of metabolites that were rescued in *Tbx1^+/-^*brains after vB12 treatment (inosine, glutamate and SCFAs). Normalized bin intensities corresponding to rescued molecules are expressed in arbitrary units. p<0.05* p<0.005** p<0.0005***

To shed light on the impact of vB12 treatment on the transcriptional profile of brain tissue in *Tbx1* haploinsufficient mice, we performed RNA quanTiseq analysis on the controlateral brain hemispheres to those used for NMR (PBS-treated *Tbx1^+/-^* and WT mice and vB12-treated *Tbx1*^+/-^ mice). We carried out the sequencing on individual hemispheres, thus each dataset was treated as a biological replicate (n = 5 for each genotype and treatment group). A principal component analysis (PCA) of gene expression revealed a similar effect in PBS-treated *Tbx1*^+/-^ and WT animals which cluster together, indicating similar transcription profiles. On the contrary, vB12-treated *Tbx1*^+/-^ animals were distant from PBS-treated *Tbx1*^+/-^ and WT animals (Figure 5B), suggesting that vB12 has a strong effect, as we observed in the metabolomics analysis. Indeed, Comparison of *Tbx1*^+/-^ animals treated with vB12 vs PBS revealed 1777 DEGs (Supplementary Table 2), of which 947 were downregulated and 834 upregulated. A pathway analysis using the KEGG database showed that vB12 treatment of *Tbx1*^+/-^ mice led to the downregulation of genes involved in neurodegeneration disorders such as Prion disease and Alzheimer disease, oxidative phosphorylation, thermogenesis and fatty acid metabolism, and the upregulation of genes involved in calcium signalling and axon guidance (Fig. 5 D and Supplementary Tab. 2).

**Figure 5.**
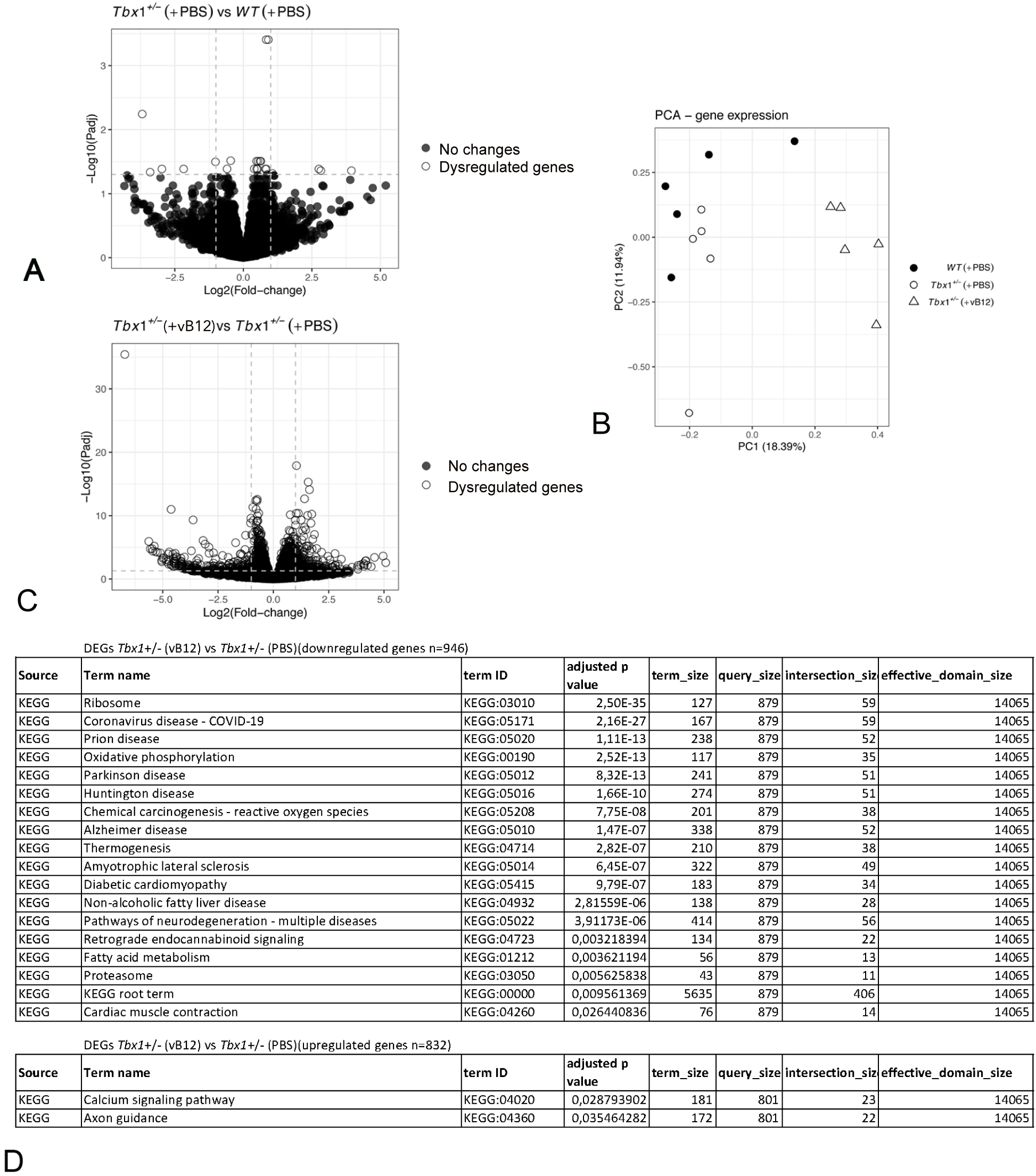
Vitamin B12 treatment has a genotype specific impact on the brains transcriptional profile. A) Gene expression changes in brain extracts of *Tbx1*^+/-^ vs WT mice treated with PBS (A) or with vB12 (C). Empty circles represent genes that were significantly dysregulated. B) A principal component analysis (PCA) plot showing similar gene expression profiles in PBS-treated *Tbx1*^+/-^ and WT mice and in vB12-treated *Tbx1*^+/-^ and WT mice, but the respective clusters are distant indicating that vB12 has a strong effect on gene expression that is independent of genotype. D) Gene ontology analysis of differentially downregulated and upregulated genes in *Tbx1^+/-^*mice after vB12 treatment.

Collectively, these data suggest that by reducing the expression of genes involved in thermogenesis and fatty acid metabolism, vB12 might slow down metabolic processes and lead to the accumulation of short chain fatty acids, thereby correcting the metabolic imbalance observed in the *Tbx1*^+/-^ brain.

### PPI deficit in *Tbx1^+/-^* mice is rescued by vB12 treatment

The consequences of the metabolic imbalance observed in *Tbx1*^+/-^ mice on brain health is unknown, but the presence of toxic metabolites such as MMA might affect brain function (Ribeiro et al., 2013)(Affonso et al., 2013). We therefore tested whether vB12 treatment was able to rescue the sensorimotor gating deficit (reduced pre-pulse inhibition, PPI) that we have reported (Paylor et al., 2006). The principle underlying the PPI paradigm is that an acoustic startle response, defined as a strong flinching response to an auditory stimulus, is attenuated when it is preceded by a weaker stimulus, the pre-pulse (Fig. 6A, centre panel). This attenuation is reduced in *Tbx1*^+/-^ mice across a range of pre-pulse intensities (Paylor et al., 2006). To test this, *Tbx1*^+/-^ and WT mice were treated postnatally with vB12 or PBS. Specifically, we treated in parallel, groups of *Tbx1*^+/-^ mice with PBS (n = 14) or vB12 (n = 13) and compared them to a group of WT mice treated with PBS (controls, n = 15). Treatment was delivered intraperitoneally twice a week for 2 months, starting at 5 weeks of age, using the dosage of 20 mg/kg, as for the other experiments described here (Fig. 6A). The last injection of vB12 or PBS was administered one day prior to testing, so that mice were off drug at the time of testing. Results showed that PBS-treated *Tbx1*^+/-^ mice had impaired PPI, as expected (Fig. 6B, Supplementary Fig. 4), whereas in vB12-treated *Tbx1*^+/-^ mice, the PPI response was indistinguishable from that of controls (Fig. 6B, Supplementary Fig. 4).

**Figure 6.**
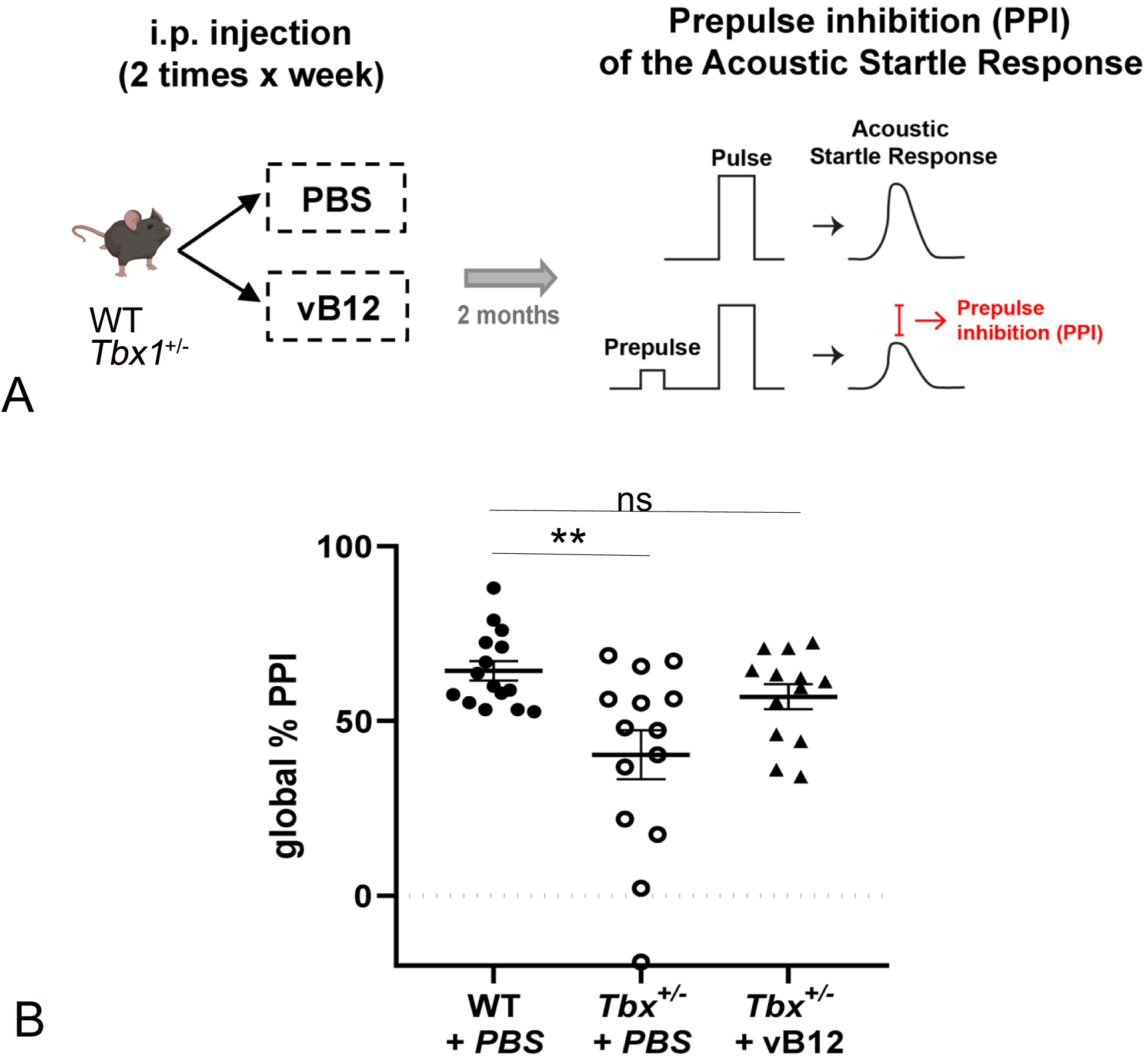
Postnatal vitamin B12 treatment rescues the PPI impairment in *Tbx1^+/-^* mice. A. Schematic representation of the experimental procedure. **B.** Dot plots showing global % PPI in WT (n = 15) and *Tbx1^+/-^*(n = 14) mice treated with PBS, and in *Tbx1^+/-^* mice (n = 13) treated with vB12. Reduced PPI in *Tbx1^+/-^* mice was rescued by vB12 treatment. ** p < 0.003 (Bonferroni’s post-hoc test).

Reduced PPI is believed to be the result of deficits in the neuronal circuitry involved in sensorimotor gating, a pre-attentive brain function that facilitates the integration of motor and sensory stimuli (Paylor et al., 2006). Our data show that prolonged postnatal vB12 treatment is able to rescue abnormal PPI responses in *Tbx1^+/-^* mice to the level of WT mice.

## Discussion

Our data show for the first time that *Tbx1* haploinsufficiency alters brain metabolism in mice. The most striking results obtained in this study were the accumulation of methylmalonic acid (MMA) in the brain of two mouse models of 22q11.2DS and the rescue of this anomaly by postnatal treatment with high doses of vB12. MMA is produced by impaired catabolism of odd-chain fatty acids, cholesterol, valine, methionine, isoleucine and threonine, with subsequent disruption of propionyl-CoA metabolism in the mitochondrial matrix and mitochondrial damage. Therefore, in order to have a broader picture of the brain metabolic profile in *Tbx1*^+/-^ mice, we performed an unbiased metabolic analysis, in addition to the preliminary targeted (candidate metabolite) study. This revealed reduced levels of leucine and increased levels of short chain fatty acids (SCFAs). Catabolism of leucine produces acetyl-CoA, which acts as a primer for carbon chain elongation of fatty acids. Moreover, we observed that the treatment of *Tbx1*^+/-^ mice with vB12, reduced MMA and restored SCFA concentrations to wild type levels. Together, these data suggest that fatty acid synthesis and mitochondrial activity are impaired in the *Tbx1*^+/-^ brain. Interestingly, TBX1 has already been linked to metabolism in adipose tissue, where its tissue specific deletion in mice impaired the glycolytic pathway and fatty acid metabolism (Markan et al., 2020).

MMA is a toxic metabolite that impacts mitochondrial activity by impairing cell respiration and glutamate uptake (Melo et al., 2012). The consequence of the latter is to increase the concentration of extracellular glutamate (Costa et al., 2022). In *Tbx1^+/-^* brain, in addition to increased MMA, we found several metabolites related to the glutamate pathway and to mitochondrial activity to be dysregulated, including reduced levels of glutamine and adenosine and increased levels of glutamate and inosine. Glutamine is a necessary metabolite for the biosynthetic process involved in purine synthesis (adenosine and inosine) (Garcia-Gil et al., 2018). Purine biosynthesis is also related to one-carbon metabolism, which generate different outputs required for nucleotide biosynthesis, as well as for the maintenance of the redox (Shuvalov et al., 2017). Moreover, purines play a role in neurotransmission and neurodevelopment (Huang et al., 2021). Therefore, the fact that vB12 treatment of *Tbx1*^+/-^ mice restored glutamate and inosine to WT levels is consistent with their metabolic relatedness and their dependence upon mitochondrial activity, which is responsive to vB12. vB12 supplementation has been shown to promote mitochondrial metabolism in human and mouse ileal epithelial cells (Ge et al., 2022).

Patients affected by methyl malonic aciduria show cognitive impairment, and in mice, MMA accumulation in the brain activates neuroinflammation and oxidative stress injury (Li et al., 2017). An additional observation is that in normal physiological conditions, glutamate is maintained at low levels in the brain. It has been demonstrated that excess glutamate affects behaviour in mice (Lander et al., 2019). Indeed, massive release of glutamate can lead to excitotoxic brain damage in mice (Lewerenz and Maher, 2015). While glutamine and glutamate were both reduced in *Tbx1*^+/-^ mice, concentrations of the inhibitory neurotransmitter GABA only increased after vB12 treatment (Supplementary Fig. 3). Hence, after vB12 treatment, increased GABA, together with the restoration of normal (low) levels of glutamate, might counteract the potentially neurotoxic excitatory effects of increased glutamate in the *Tbx1^+/-^* brain.

Another important finding of our study was the discovery that postnatal vB12 treatment of *Tbx1*^+/-^ mice rescued the reduced PPI phenotype first reported by Paylor et al. in 2006. The mechanisms by which *Tbx1* mutation affects brain development and brain function are unknown, but they are cell non-autonomous mechanisms (Flore et al.,, 2017) because in the mouse brain, *Tbx1* expression is limited to brain vessels, and it is not expressed in neurons or other brain cell types (Paylor et al., 2006)(Cioffi et al., 2014). In mice, *Tbx1* loss of function causes hyperbranching and disorganization of the brain vascular network and tissue hypoxia (Cioffi et al., 2014), all of which might alter brain metabolism and behaviour. Thus, the phenotypic rescue might occur at the metabolic or vascular level, but given that it was obtained when vB12 treatment was started in juvenile mice, when the brain vascular network is fully formed, we favor the metabolic route. In addition, the observed upregulation of axon guidance genes might enhance sensorimotor function.

Although, transcriptional analysis performed here revealed only 22 DEGs between *Tbx1^+/-^* and wild type brains and 3 of them are involved in mitochondrial activity, further studies will be necessary to assess their role in the behavioural phenotype. The reason for the low number of DEGs might be technical rather than biological, i.e., in the nature of bulk RNA-seq, which is not very sensitive to changes in gene expression in cell populations that are not well represented in the tissue analysed. In the adult mouse brain, *Tbx1* is only expressed in a subpopulation of brain vessels (Paylor et al., 2006)(Cioffi et al., 2014).

Could impaired behavioural phenotype in *Tbx1^+/-^* mice be due to the increased levels of glutamate and glutamine reported here? A recent study reported that 22q11DS patients with psychotic symptoms had increased levels of glutamate and glutamine in the hippocampus and in the superior temporal cortex (Mancini et al., 2023). Previously, increased glutamate and myo-inositol were found in the hippocampus of 22q11DS patients with schizophrenia (da Silva Alves et al., 2011). Future studies will be needed to elucidate the role of these metabolites in the behavioural anomalies in the 22q11DS mouse models.

In summary, our study shows for the first time that in adult mice, *Tbx1* haploinsufficiency alters brain metabolic homeostasis and that this is rescued by postnatal vB12 treatment. Additionally, we report that the same pharmacological treatment rescues PPI anomalies in *Tbx1* mutants, suggesting that they are related to the underlying metabolic changes caused by the mutation. Definition of the molecular mechanisms that lead to these metabolic anomalies will require further studies. Nevertheless, our data open a new direction for the design of a therapeutic strategy that could be applied postnatally and might help to combat the brain-related phenotype of 22q11.2DS.

## Material and Methods

Animal research was conducted according to EU and Italian regulations. The animal protocol has been approved by the animal welfare committee of the Institute of Genetics and Biophysics (Organismo per il Benessere Animale, OPBA-IGB), protocol n. 7E58D.24. We used the following mouse lines in C57/Bl6N background; *Tbx1 ^lacZ^*^/+^, *Tbx1^ΔE5/+^*, *Df1*/+ adult mice (Lindsay et al., 2001)(Xu et al., 2005).

### Mouse treatment

*Tbx1^+/-^* and WT mice were injected intraperitoneally with vB12 dissolved in PBS 20 mg/kg (stored away from light) or with PBS, twice a week for 4 weeks starting at 4 weeks of age for all experimental procedures. In addition, for behavioural analysis, all animals were treated until the day before behavioural testing that was performed at 12-13 weeks of age. For metabolomic and molecular studies, at the end of treatment (at 12 weeks of age), the brains were surgically removed after cervical dislocation and stored at –80 °C

### Metabolomic analysis of mouse brains LC-MS/MS

Metabolites were extracted from mice brain tissues (*Tbx1^+/-^* and WT) for targeted metabolome analysis by liquid chromatography - tandem mass spectrometry (LC-MS/MS). Frozen tissues were disrupted in 50:50 cold methanol/0.1 M hydrogen chloride and homogenized at high speed by shaking with stainless-steel beads into a TissueLyser LT (Qiagen, Duesseldorf, Germany). The metabolites-containing supernatant was isolated from the protein pellet by centrifugation at 13,000 rpm for 60 min at 4°C. Proteins were resuspended and quantified for the subsequent normalization data as published (Gonzalez Melo et al., 2021a). The metabolite extracts were alkalinized to pH 7– 8 and dried under nitrogen. Finally, dried metabolite mixtures were resuspended in methanol and analysed by targeted MS/MS to determine the contents of amino acids (AAs), acylcarnitines (ACs), methylmalonic acid and lactic acid. The metabolomic platform was developed to perform the fast identification of biomarkers of inherited metabolic diseases and adapted for the purpose to the present analytical work. Prior to MS analysis, a standard mixture containing labelled AAs and ACs was added to all of the samples for the derivatization of all the endogenous and exogenous molecules, as previously reported (Giacco et al., 2019). Finally, the MS/MS experiments were carried out in flow injection analysis (FIA) mode for AAs and ACs and in LC for methylmalonic acid (MMA) and lactic acid (LA) by using an API 4000 triple quadrupole mass spectrometer (Sciex, Toronto, Canada) coupled with an 1160 series Agilent high-performance liquid chromatography (HPLC) system (Agilent Technologies,Waldbronn, Germany) (Gonzalez Melo et al., 2021b)(Campesi et al., 2023).

The chromatographic separation of MMA and LA was performed with a 3 µm analytical column Phenomenex Gemini C6-Phenyl 100×2.0 mm (Phenomenex, Torrance, CA, USA). Solvent A and B were consisted of 0.1% formic acid in water and acetonitrile, respectively. The flow was set at 300 µL/min with the following gradient: 0.0-0.5 min: 30% B; 0.5-3.0min: 30% to 90% B; 3.0-3.10 min: 90% to 30% B; 3.1-4.0 min: 30%. MRM experimental parameters were: polarity: negative; Q1/Q3 (m/z): 117.00/73.00 (MMA), 89.00/59.00 (LA); DP (volts): –34(MMA), –50 (LA); CE (volts): – 12 (MMA), –14 (LA). Quantitative analysis of the data was performed with ChemoView v2.0.4 software through the comparison of the analytes and their corresponding internal standard areas.

### Multivariate statistical data analysis of LC-MS/MS data

Univariate statistics data analysis was carried out with GraphPad Prism 9.0 and the results are presented as the mean ± standard error of the mean (SEM). The data were normalized to convert values from different data sets to a common scale, defining zero as the smallest value and one hundred as the largest value in each data set. The normalized metabolic data set was log10-transformed. The volcano plot was performed to visualize the predictive component loading and identify significantly altered metabolites by their content variation (Difference) and statistical significance –log10 p-value). The normalized data set was then used to perform pattern correlation analyses with MetaboAnalyst 5.0 (http://www.metaboanalyst.ca).The statistical significance of the difference in metabolite samples concentrations between two different groups was evaluated by parametric (unpaired t-test with Welch correction) or non-parametric (Mann–Whitney test) comparisons. The significant difference between multiple groups was evaluated by ordinary one-way ANOVA and Hold–Sidak’s multiple comparison test in normally distributed data sets, and Kruskal–Wallis test and Dunn’s multiple comparison test in non-normally distributed data sets. The normal distribution was verified according to the D’Agostino and Pearson test.

### NMR

Combined extraction of polar and lipophilic metabolites were carried out by using methanol/water/chloroform as suggested by the Standard Metabolic Reporting Structures working group (Lindon et al., 2005). Standard metabolic reporting structures working group. Summary recommendations for standardization and reporting of metabolic analyses. Polar and nonpolar fractions were then transferred into glass vials and the solvents removed by using a rotary vacuum evaporator at room temperature and stored at -80°C until they were analysed. For NMR analysis, polar fractions were resuspended in 630 µl of phosphate buffer saline (PBS, pH 7.4), adding 70 µl of ^2^H_2_O solution [containing 1 mM sodium 3-trimethylsylyl [2,2,3,3-2H4] propionate (TSP) as a chemical shift reference for ^1^H spectra] to provide a field frequency lock, reaching 700 µl of total volume. The nonpolar fractions were resuspended in 700 µl of C^2^HCl_3_. Samples were loaded into the autosampler and NMR spectra and acquired on a Bruker Avance III–600 MHz spectrometer (BrukerBioSpin GmbH, Rheinstetten, Germany), equipped with a TCI CryoProbe fitted with a gradient along the Z-axis, at a probe temperature of 27°C. In particular, standard 1D proton spectra and 2D experiments (clean total-correlation spectroscopy TOCSY and heteronuclear single quantum coherence HSQC) were acquired providing monodimensional metabolic profiles and homonuclear and heteronuclear spectra for metabolite identification. Metabolite assignments were achieved by comparing signal chemical shift with literature and online databases. All 1D spectra were processed and automatically data reduced in bins and arranged as a data matrix with AMIX 3.9.7 package (Bruker Biospin GmbH, Rheinstetten, Germany) then imported into the SIMCA14 package (Umetrics, Umeå, Sweden) for multivariate data analysis.

### Multivariate statistical data analysis of NMR data

To discriminate between treated and untreated mice in response to VitB12 administration and according to their brain NMR profiles, multivariate statistical analysis was performed. In particular, the unsupervised principal component analysis (PCA) was first applied to assess class homogeneity, uncover data trends and detect outliers. Then, supervised methods such as Orthogonal Partial Least Squares Discriminant Analysis (OPLS-DA) were used to improve class separation, thus better appreciating clusters and the spectral variables influencing samples distribution according to the alteration of their metabolic profile. The performance of each supervised model was estimated by evaluating the percent of data variation explained (R^2^) and the one predicted by according to cross validation (Q2). Moreover, OPLS-DA models were validated by internal iterative cross-validation with 7 rounds permutation test response (800 repeats), and CV-ANOVA (ANOVA testing of Cross-Validated predictive residuals). Data visualization was achieved through scores, loadings and S plots, highlighting specific compounds as putative markers. To identify the subset of most responsible metabolites characterizing class discrimination, NMR variables were selected considering the correlation loadings values |p(corr)|> 0.7 (as shown in the S-plots in the supplementary figures) and tested them for univariate statistical analysis using unpaired t-test for two class analysis (once considered variance homogeneity and Welch correction) or ANOVA test with Bonferroni correction in case of multiple comparisons, after assessing gaussian distribution with normality test (Shapiro-Wilk). Statistical tests were elaborated with the OriginPro 9.1 software package (OriginLab Corporation, Northampton, USA) and R software [R core team (https://www.r-project.org/).

### Pathway analysis

Pathway topology and biomarker analysis on selected and more representative metabolites in class separation was applied to the pathway topology search tool in Metaboanalyst 5.0 (Pang at al 2021). The Pathway Analysis approach integrates results from both pathway enrichment analysis and the pathway topology analysis to spot the most relevant pathways involved in the study. Mouse library was chosen and analysed using Fisher’s Exact Test for over representation and Relative-betweenness Centrality for pathway topology analysis.

### RNA isolation

Total RNA was isolated from whole brains with Qiazol (QIAGEN) according to the manufacturers protocol with an additional precipitation step.

### Transcriptomic Quantiseq analysis

QuanTiseq libraries were prepared and sequenced at Telethon Institute of Genetics and Medicine (TIGEM), Naples. The facility performed quality control and trimming for each sample aligned them to the mouse genome (mm10) using STAR (Dobin et al., 2013), and provided the gene expression as a raw count matrix. We imported the count matrix in the R environment, evaluated the counts per million (CPM) expression, and retained in our subsequent analysis only those genes with CPM > 1 (as expressed genes) in at least three samples. We used the EdgeR package (McCarthy et al., 2012), i.e., first, we estimated the library size using the calcNormFactors function. Then, we defined the design matrix, with the three experimental conditions and the corresponding biological replicates, estimating the parameters of the negative binomial model using the estimateGLMRobustDisp function. Finally, we fitted the model with the glmFit function and extracted the contrast of interest from the joint fit. We used the glmLRT function to test the differential expression and we applied the decideTests function with method = “separate” and FDR<0.05 for each contrast of interest. Pathway analysis was performed by using the gProfiler online tool. In all condition, the lists of differentially expressed genes were analysed, using all expressed genes as gene background.

### Behavioural analysis

#### PPI – Acoustic startle response and pre-pulse inhibition

The test was performed using the SR-LAB^TM^ Startle Response System apparatus (San Diego Instruments, CA, USA), as previously described (Mandillo et al., 2008)(Mandillo et al., 2013). Briefly, the PPI session started with a 5 min habituation period, in which only a background noise (BN) of 72-73dB was present, followed by 5 consecutive startle trials which were excluded from the statistical analysis and 9 different trial types repeated 10 times in random order with an inter-trial interval of 25 s on average. The startle trial consisted of the presentation of a 110dB/40ms acoustic pulse (white noise); the “no stimulus” trial consisted of BN/10ms presentation to measure the baseline movement of the mouse; the 4 pre-pulse alone trials consisted of the presentation of 10ms long 76, 80, 88 or 100dB stimuli; the 3 pre-pulse inhibition (PPI) trials consisted of 10ms long stimuli of 76, 80 or 88dB followed after 50ms by a 110B/40ms pulse. For each trial, the startle response was recorded every millisecond for 65ms following the onset of the acoustic pulse. Maximal peak-to-peak amplitude was used to determine the ASR in the acoustic startle pulse and/or prepulse alone trials. Mice were off drug during the whole procedure.

PPI was expressed as %PPI = [100 - (startle response for prepulse/startle response for startle-alone trials) x 100]. The whole PPI session lasted about 45 min. Average % PPI (global % PPI) over the sampling period was taken as the dependent variable (Iemolo et al., 2023).

% PPI was analysed with a two-way ANOVA for repeated measures for the factors: prepulse (three levels % PPI 76p110, % PPI 80p110 and % PPI 88p110) and treatment (three levels: WT PBS, *Tbx1^+/-^* PBS and *Tbx1^+/-^* vB12). A Bonferroni correction was applied for global % PPI.

## Acknowledgements

We thank Rosa Ferrentino and Marchesa Bilio for expert laboratory assistance. We also thanks IGB mouse facility and especially Lucia Mele for expert technical assistance.

This work was funded by the Ministry of University and Research (MUR), National Recovery and Resilience Plan (NRRP) 2022, project MNESYS (PE0000006) to AB; PRIN-PNRR 2022, project P2022ARZ5J to GL, MR and EI; and DBA.AD005.225 NutrAge FOE 2021 to EDL.

