## Supplementary figures and images for "Tbx1 haploinsufficiency causes brain metabolic and behavioral anomalies in adult mice which are corrected by vitamin B12 treatment"

### Supplementary Figure 1

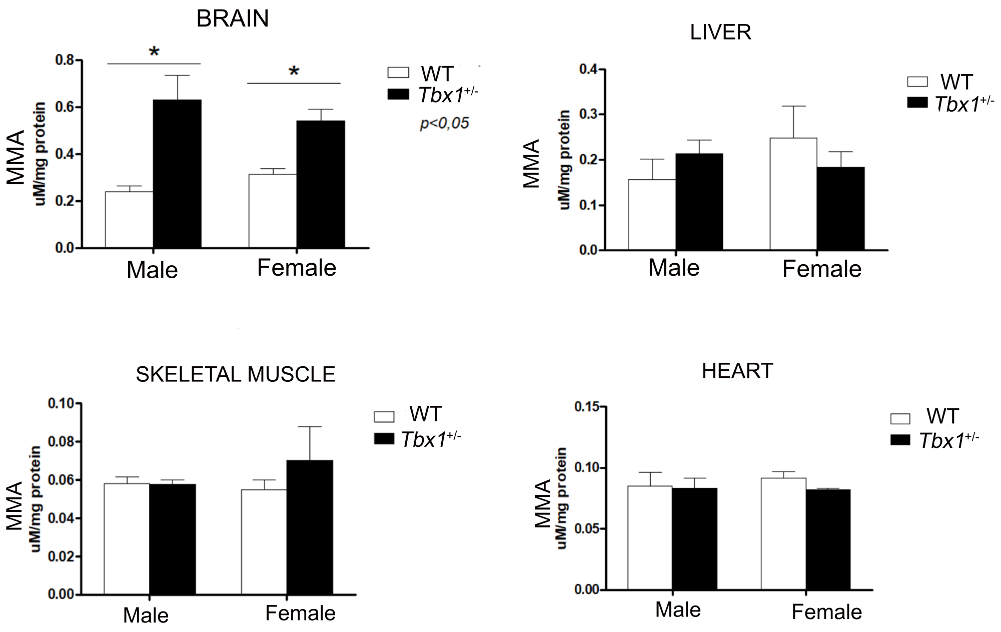


Supplementary 1

### Supplementary Figure 2

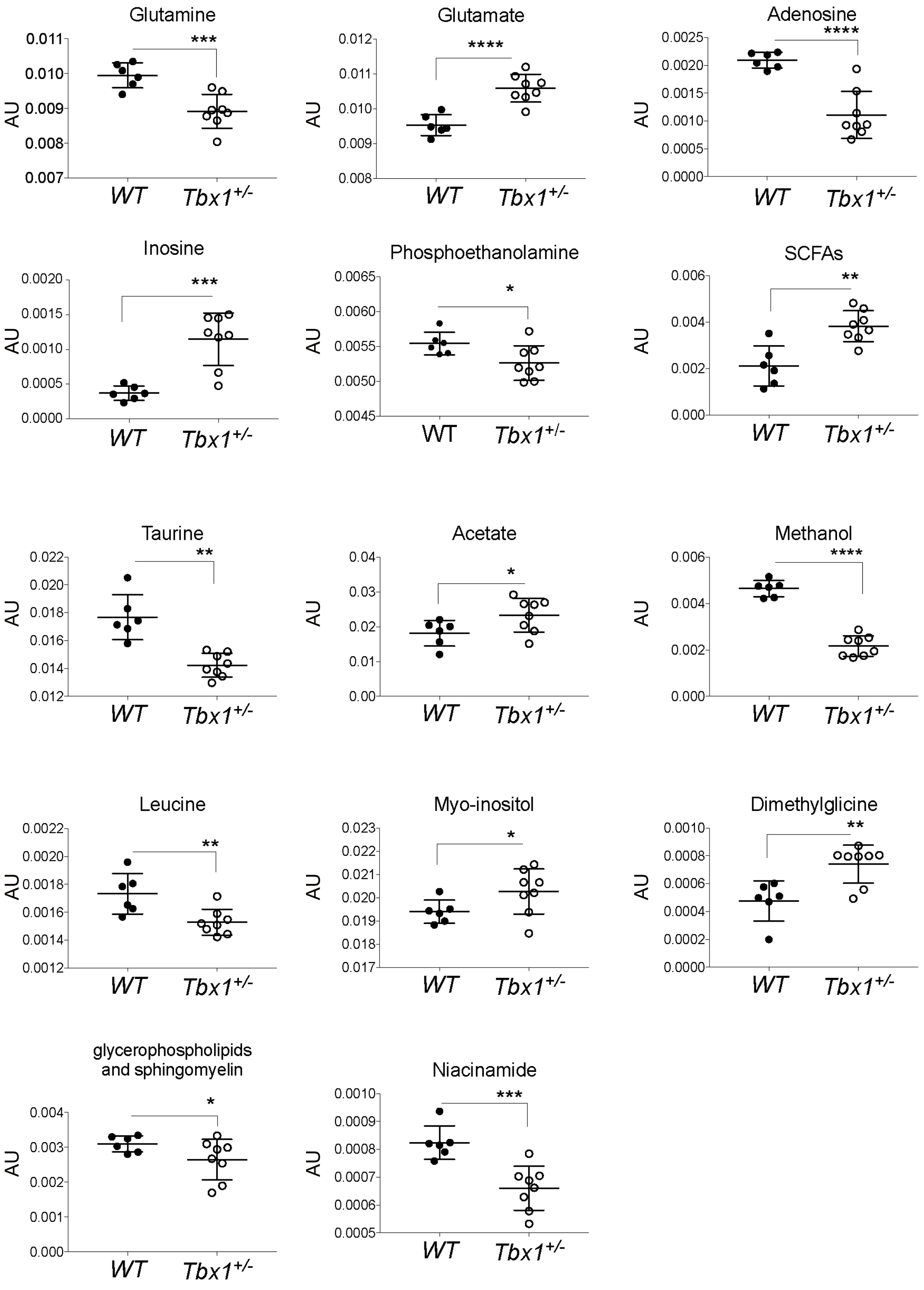


Supplementary 2

### Supplementary Figure 3

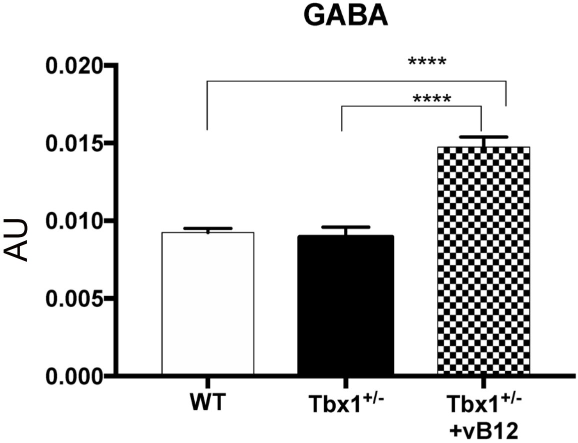


Supplementary Figure 3

### Supplementary Figure 4

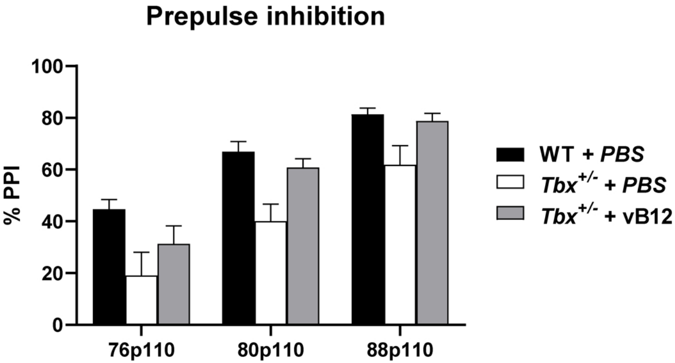


Supplementary Figure 4
